# The species richness of the *Salix viminalis* rhizosphere at the Stebnyk tailings storages is dependent on supplementation from the *Salicornia europaea* rhizosphere

**DOI:** 10.1101/2023.09.25.559290

**Authors:** Anastasiia Fetsiukh, Taavi Pall, Salme Timmusk

**Author notes:** Equal contribution: Co-first (Anastasiia Fetsiukh, Taavi Pall).

## Abstract

Manipulating the rhizosphere microbiome to enhance plant stress tolerance is an environmentally friendly technology and a renewable resource to restore degraded environments. Here we considered the *Salicornia europaea* rhizosphere community, and the ability of the phytoremediation plant *Salix viminalis* to recruit its beneficial microbiome to mediate the pollution stress at the Stebnyk mine tailings storage. The tailings contain large amounts of brine salts and heavy metals that contaminate the ground water and surrounding areas, changing soil biogeochemistry and causing increased erosion. The species richness of the endophytic bacterial community of *S. viminalis* roots was assessed based on observed OTUs, Shannon-InvSimpson, and evenness index. Our results show that biodiversity was decreased across the contamination zones and that *S. europaea* supplementation significantly increased the species richness. Our results also indicate that the number of dominating OTUs was not changed across zones in both *S. europaea*-treated and untreated bacterial populations, and that the decrease in richness was mainly caused by the low abundance of OTUs.

The importance of engineering microbial communities that support the genetic diversity of degraded environments and the challenges with high throughput metabarcoding databases are discussed.

## 1. Introduction

Plant-soil-microbe interactions play an important role in plant community performance since microbial abundance and composition largely determine the health and productivity of any soil (Hartmann, Frey et al. 2015, Timmusk, Behers et al. 2017, Averill, Anthony et al. 2022). Plant growth-promoting microorganisms form beneficial relationships with plants, and their interactions evoke local and systemic responses in plants through several molecular and cellular mechanisms upon perception of environmental signals. This enables plants to cope with stresses such as temperature extremes, drought, flooding, salinity, and heavy metal pollutants (Timmusk, Behers et al. 2017, Timmusk, Pall et al. 2023). Hence, the ability to manage the soil microbiome under pollution stress is likely to be the key to efficient bioremediation (Batista and Singh 2021). Since rhizobacteria were first described (Hiltner 1904), single microbial isolates as well as microbial consortia have been used over decades for inoculation to influence the host plants to remediate various stress situations (Bashan 1998, Timmusk and Wagner 1999, de-Bashan, Hernandez et al. 2012, Glick 2012, Timmusk, Abd El-Daim et al. 2014, Ray, Lakshmanan et al. 2020). Environmental remediation using microbial inocula has several advantages over artificial chemical treatments (Sonawane, Rai et al. 2022, Tufail, Iltaf et al. 2022). As in natural conditions 95% of microbes are bacteria, plant growth promoting rhizobacteria (PGPR) have a special potential for bioremediation (Santoyo, Urtis-Flores et al. 2021, Orozco-Mosqueda, Fadiji et al. 2022). The rhizosphere microbiome is an environmentally friendly renewable resource, and activation of the resource restores soil and plant health (Bar-On, Phillips et al. 2018, Timmusk, Conrad et al. 2020, Averill, Anthony et al. 2022). At the same time there are general challenges that hinder the breakthrough of microbial inocula into commerce; in particular, inconsistent colonization and inefficiency of field application (Timmusk, Behers et al. 2017, Dini-Andreote and Raaijmakers 2018, Oyserman, Medema et al. 2018, Timmusk, Seisenbaeva et al. 2018, Kaminsky, Trexler et al. 2019, Ray, Lakshmanan et al. 2020, Trivedi, Leach et al. 2020, Trivedi, Mattupalli et al. 2021, Timmusk and de-Bashan 2022, Timmusk, Pall et al. 2023). Historically PGPR strains have been selected based on their biochemical properties (Timmusk, Behers et al. 2017, Ray, Lakshmanan et al. 2020). It is however highly important to consider microbial communities and plant complementary traits from the indigenous communities in a holistic manner (Timmusk, Behers et al. 2017, Ray, Lakshmanan et al. 2020, Timmusk, Pall et al. 2023). Here we considered the principle that while the bacterial rhizosphere microbiome is rather dynamic and unstable, there are endophytic communities that present relatively stable communities (Timmusk, Behers et al. 2017, Timmusk, Pall et al. 2023). The other principle we considered is host-mediated microbiome selection (HMMS): a strategy that focuses on the host organism’s intrinsic ability to recruit its own beneficial microbiome (Baldassarre, Ying et al. 2022, Delannoy-Bruno, Desai et al. 2022). The microbiome can then be characterized by the high throughput metabarcoding (HTM) approach. (Timmusk, Behers et al. 2017, Dini-Andreote and Raaijmakers 2018, Oyserman, Medema et al. 2018, Timmusk, Seisenbaeva et al. 2018, Kaminsky, Trexler et al. 2019, Ray, Lakshmanan et al. 2020,Timmusk and de-Bashan 2022).

One of the examples where bioremediation is greatly needed is a tailings storage on the north-eastern outskirts of the Stebnyk (Lviv region, Ukraine), which contains 22 million tons of waste including clay material, undissolved salts, brine with a high content of sodium chloride, heavy metals (HM) and salts (Fetsiukh, Bunio et al. 2022). The waste causes salinization of groundwater, reservoirs, and surrounding areas (Fetsiukh, Bunio et al. 2022). The negative effects of salt pollution include changes in soil biogeochemistry and composition, and characteristics such as decreased aeration and decreased soil permeability with increased erosion. Currently developed hydro-technical technologies at the tailing storage have to be followed by carefully planned bioremediation strategies (Mokryi, Petrushka et al. 2023). Here we considered the potential of the basket willow (*Salix viminalis*) for the tailing storage bioremediation. HTM has advanced our understanding of microbial diversity, distribution and function as well as the ecological roles of members of the community (Bahram, Hildebrand et al. 2018, Oyserman, Medema et al. 2018)(de Vries, Griffiths et al. 2018) This method allows microbial taxa to be grouped into ecologically meaningful molecular operational units that represent taxonomic groups OTUs; (Taberlet, Coissac et al. 2012, Alivisatos, Blaser et al. 2015, Hultman, Waldrop et al. 2015, Gilbert, Quinn et al. 2016, Jansson and Hofmockel 2018, Jansson and Hofmockel 2020, Naylor, McClure et al. 2022). A major challenge with bacterial taxonomy and OTUs is, however, the fluid nature of their genomes (Prosser, Bohannan et al. 2007). The databases of bacterial taxonomy do not consider the gene-transfer processes of bacteria. Genomic processes in rhizosphere can transfer a small part of the genome that replaces the homologous copy in the genome, and maintains the cohesion of the species (Timmusk, Pall et al. 2023). Alternatively, gene transfer processes can result in the transfer of genes that have no counterparts in the bacterium that receives them. This DNA can be maintained on a plasmid, or integrated into the host genome by non-homologous recombination (Timmusk, Pall et al. 2023). A result of gene transfer can be that the bacterial genome contains two distinct parts, the core genome comprising genes that are normally essential and an accessory genome providing special ecological adaptations from genes that are easily gained or lost (Prosser, Bohannan et al. 2007). Strains of the same species, as defined by their core genome, can differ with respect to hundreds of accessory genes, and have different ecological potentials (Prosser, Bohannan et al. 2007). Studies of 16S ribosomal RNA (rRNA) gene sequences show the vast diversity of bacterial communities, but if the accessory genome confers much of the important ecological adaptation then the true ecological diversity depends on the rich mixture of catabolic plasmids, resistance transposons and genetic islands (Prosser, Bohannan et al. 2007) which with higher probability are gained from the harsh habitats (Timmusk, Paalme et al. 2011, Timmusk, Abd El-Daim et al. 2014, Timmusk, Pall et al. 2023). The accessory genome can be shared among unrelated bacteria in an environment that favors them but can be absent in the ‘same’ bacterial species growing elsewhere (Timmusk, Paalme et al. 2011) Timmusk, Pall et al. 2023). This ability to pick up genetic information is mostly linked to bacteria from harsh environments and is used here as a third principle; and the rhizosphere of *Salicornia europaea* at Stebnyk was applied as a genetic resource for bioremediation. The *S. europaea* plants were grown under the salt and HM pollution stress over four decades. Former research shows that *Salicornia* species may provide a microbiome that effectively attenuates saline stress (Abdollahzadeh, Niazi et al. 2019, Hrynkiewicz, Patz et al. 2019). Here we hypothesized that bacteria that have been living under HM and salt pollution stress over a long period of time have gained the potential to mitigate plant pollution stress. Hence as a first step we determined the pattern of endophytic bacterial colonization of *S. viminalis* across the Stebnyk contamination gradient. We predicted that *S. europaea* treatments would increase the abundance and species richness of the endophytic community. We tested these hypotheses by conducting DNA metabarcoding along replicated pollution zones.

## 2. Material and Methods

### 2.1. Sampling and experimental design

In total, thirty-six (36) individual cuttings were examined at a gradient of Stebnyk mine tailing representing three contamination levels earlier described (Fetsiukh, Bunio et al. 2022) (Fig 1). In this project the investigations were carried out in three regions that were weakly, moderately and heavily contaminated (Zone 1, 2 and 3 respectively) (Fig 1). Samples were taken at a depth of 0–25 cm at places of strong (49°18’45.2”N 23°34’05.5”E; 49°18’45.4”N 23°34’03.4”E; 49°18’45.3”N 23°34’04.5”E) and medium salinity (49°18’45.0”N 23°34’07.7”E; 49°18’43.8”N 23°34’07.8”E; 49°18’43.3”N 23°34’07.4”E), as well as in places of restored biogeocenosis (49°18’39.8”N 23°33’59.3”E; 49°18’40.0”N 23°34’00.7”E; 49°18’41.1”N 23°33’57.7”E), which was determined visually at the place of plant growth. To explore the diversity, samples were taken from six different sites at each of the three regions and used as growth substrate for willow (*Salix viminalis* L.) cuttings. The cuttings were left to grow in pots with the soils sampled at the three regions as well as with soil samples that were enriched with slurries from the *S. europaea* rhizosphere.

**Fig 1.**
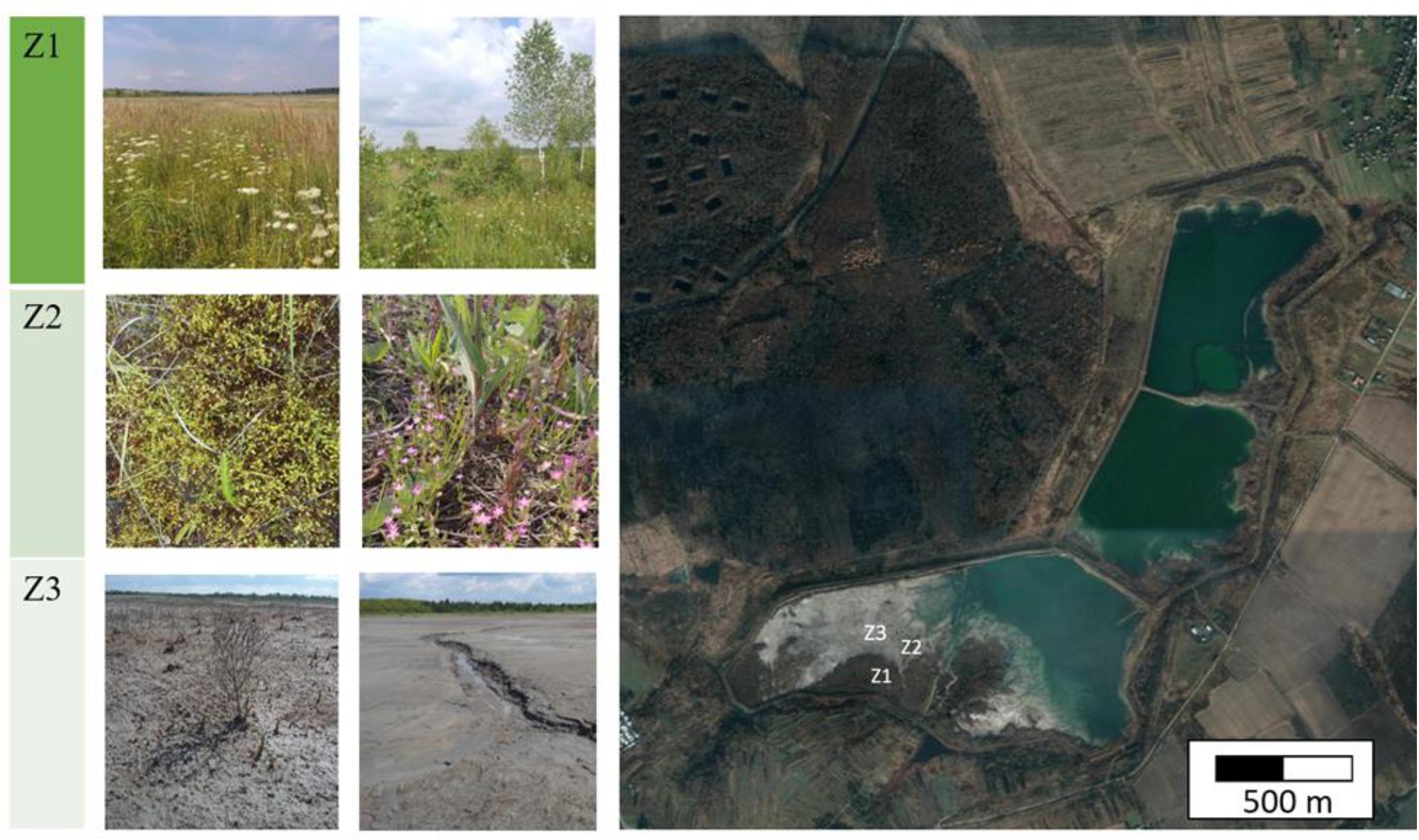
Sampling locations on the Stebnyk tailings storage: Z1 – places of restored biogeocenosis; Z2 — places of distribution of pioneer glycohalophytes (medium salinity); Z3 — places of minor distribution of *Salicornia europaea* L. (strong salinity).

The rhizosphere microbiome of *S. europaea* L. was collected on Stebnyk tailing under field conditions and transferred to sterile bags, which were transported to the laboratory.

The rooted cuttings of *S. viminalis* were planted in pots according to the following scheme: 5 cuttings in each pot that contained 2.5–3 kg of tailing’s soil (Z1, Z2, Z3); 5 cuttings in each pot that contained 2.5–3 kg of tailing’s soil on the bottom and upper layers, and a middle layer of potting soil with native bacteria *Salicornia europaea* L. (Z1plus, Z2plus, Z3plus). As a control, 5 rooted cuttings were grown using a sand culture. After three months of growth the willow cuttings were harvested, and DNA extracted from the below-ground part of the plants.

### 2.2 DNA extraction and PCR amplification

DNA was extracted from the roots grown in the pots with soil from the zones 1, 2 and 3 (Z1, *Z2 and Z3*) and from the ones where the growth substrate contained soil from zones enriched with the *S. europaea* rhizosphere soil (Z1 plus, Z2 plus and Z3 plus).

Microbial DNA was obtained from 200 mg of *S. viminalis* roots using the Nucleo Spin® Soil kit (Macherey-Nagel, Germany). Quantitative determination of the concentration (ng/μl) and purity of DNA (A260/280) was performed spectrophotometrically using a NanoDrop Lite Spectrophotometer (Thermo Fisher Scientific). DNA concentration was determined fluorometrically using Qubit 2.0 (Invitrogen). The quality of the DNA was checked via 1% agarose gel electrophoresis. DNA samples were stored at -20°C for further analysis.

Prokaryotic 16S rRNA gene fragments were amplified by a two-step PCR procedure where the first step consisted of 2.5 ng extracted DNA, 2X Phusion PCR Mastermix (Thermo Scientific, Waltham, MA, US) and 10 μM of the primers pro341F/pro805R in 15 μl reactions. Two independent PCRs were run under the following conditions: 3 min at 98 °C, followed by 25 cycles of 98°C for 30 s, 55°C for 30 s and 72°C for 30 s and a final extension step of 10 min at 72°C. The PCR products were then pooled and checked by 1% agarose gel electrophoresis. A single 30 μl reaction was performed for the second PCR, using 2 μM of primers with Nextera adaptor and index sequences, and 3 μl of the pooled PCR product from the first PCR. Conditions were the same as the first step, except for an annealing temperature of 55°C and an extension time of 45 s, with 8 cycles. The final PCR products were purified using an E.Z.N.A.® Cycle-Pure Kit (Omega Bio-tek, Georgia) following the manufacturer’s instructions. The amplicon size was checked by gel electrophoresis and the quality control was performed on a BioAnalyzer (Agilent, Santa Clara, CA, US). After quantification using a Qubit fluorometer (Invitrogen, Carlsbad, CA, US), libraries were created. Sequencing was performed on an Illumina MiSeq instrument using the 2,250 bp chemistry.

### 2.3. Processing of the reads, clustering and taxonomic identification

16S rRNA analysis of MiSeq sequencing data was performed essentially as described in mothur MiSeq SOP (Kozich, Westcott et al. 2013). Paired end reads were merged with make.contigs command using parameters “align=needleman, match=1, mismatch=-1, gapopen=-2, gapextend=-1”. Mean alignment length was 466 bases, 97.5% sequences had 0 ambiguous bases. Merged reads were trimmed using screen.seqs command with “maxlength=464, maxambig=0” parameters. Duplicate sequences were removed using unique.seqs command and aligned to custom silva.v3v4.align reference alignment using align.seqs command. Custom 16S rRNA v3v4 region reference alignment was generated using *L. acidophilus* 16S rRNA gene sequence (EF533992.1) and SILVA seed reference file (silva.seed_v123.align, https://mothur.s3.us-east-2.amazonaws.com/wiki/silva.seed_v123.tgz). Briefly, the *L. acidophilus* 16S rRNA sequence covering the region amplified by pro341F/pro805R oligos was aligned to the SILVA reference file with mothur align.seqs command to obtain coordinates for trimming the SILVA reference. Then, the SILVA reference was trimmed to the v3v4 region using start and end coordinates 6388 and 25316, respectively (465 bases), with the mothur pcr.seqs command. After alignment to reference, poorly aligned sequences and sequences with long homopolymer stretches were removed using the screen.seqs command with parameters start=1, end=18928, maxhomop=8. Overhangs on either side of the v3v4 region were removed using the filter.seqs command with parameters vertical=“yes” and trump=“.”Almost identical sequences with maximum 4 differences were merged using the pre.cluster command with the diffs=4 parameter. Chimeric sequences were identified and removed from further analysis using the chimera.vsearch and the remove.seqs command. Retained sequences were classified taxonomically using the classify.seqs command with reference=“trainset16_022016.rdp.fasta” and taxonomy=“trainset16_022016.rdp.tax”, where v16 RDP reference files were obtained from mothur (https://mothur.org/wiki/rdp_reference_files/). Sequences classified as belonging to taxa “Chloroplast-Mitochondria-unknown-Archaea-Eukaryota” were removed from further analysis. Sequences were clustered to OTU-s using the cluster.split command with parameters taxlevel=4 and cutoff=0.03. OTU counts were obtained using command make.shared, and the consensus taxonomy of each OTU was obtained using the classify.otu command. The mothurs’ make.biom command was used with inputs from make.shared and classify.otu to generate a biom v1.0 file for downstream analyses.

### 2.4. Statistical analysis

For Bayesian modelling we used the R libraries rstan vers. 2.21.3 (Stan Development Team 2020) and brms vers. 2.16.1 (Burkner, Doebler et al. 2017). Models were specified using extended R lme4 (Bates et al. 2015) formula syntax as implemented in R brms package. We used weak priors to fit models. We ran minimally 2000 iterations and four chains to fit models. When suggested by brms, Stan NUTS control parameter adapt_delta was increased to 0.95– 0.99 and max tree depth to 12–15. Hypotheses were tested using the alpha = 0.05 level, specifying the Bayesian confidence interval (credible interval), containing 1-alpha = 0.95 (95%) of the posterior values, to determine the presence of an effect.

## 3. Results

Thirty-six (36) *S. viminalis* root samples were studied representing the 1,455,675 (after rarefaction 245,448) reads, comprising 26,611 unique OTUs (after rarefaction 8,255). A total of 12,133 (after rarefaction 4,209) and 19,731 (after rarefaction 5,917) OTUs were analyzed from the *S. viminalis* roots grown in the unsupplemented soils from Z1-3 and in the soils from Z1-3 supplemented with *S. europaea* rhizosphere, respectively. The lowest number of OTUs was found at Z3 (3,537, after rarefaction 1,311) and the highest number was at Z1 supplemented with *S. europaea* rhizosphere (9,423, after rarefaction 3,690). The roots from all three zones had their own characteristic community after rarefied data and only 323 and 341OTUs were present at all three zones presenting the cuttings grown with and without *S. europaea* rhizosphere, respectively. This amounts to 7.6%, 95% CI [6.8%, 8.4%] and 5.8%, 95% CI, [5.2%, 6.4%] of the total number of OTUs, respectively (Fig 2 A, B). Analysis suggests that the plants grown in soils supplemented with *S. europaea* rhizosphere had 1.9 (95% CI [0.9, 2.8]) percentage points fewer OTUs common to all three zones than plants grown in unsupplemented soils, the probability of a non-zero effect was nearly 1. Accordingly, there were more unique OTUs in samples supplemented with *S. europaea* rhizosphere than in unsupplemented samples (Fig 2 C). Analysis of pairs of unsupplemented versus *S. europaea* rhizosphere-supplemented OTU sets at three zones showed that there was a higher proportion of unique OTUs relative to common OTUs in cuttings grown in soils supplemented with *S. europaea* rhizosphere from Z1 and Z3 (Fig 2 D, F, G). Whereas, in Z2 there were less unique OTUs relative to common OTUs in plants grown in soils supplemented with *S. europaea* rhizosphere. (Fig 2 E, G). The number of OTUs common between plants grown either in *S. europaea* rhizosphere supplemented or unsupplemented soils decreased with increasing contamination from Z1 to Z3 (Fig 2 D-F). 203 OTUs out of 8,255, comprising 2.5%, 95% CI [2.1%,2.8%] of total OTUs, were present in plants from all conditions.

**Fig 2.**
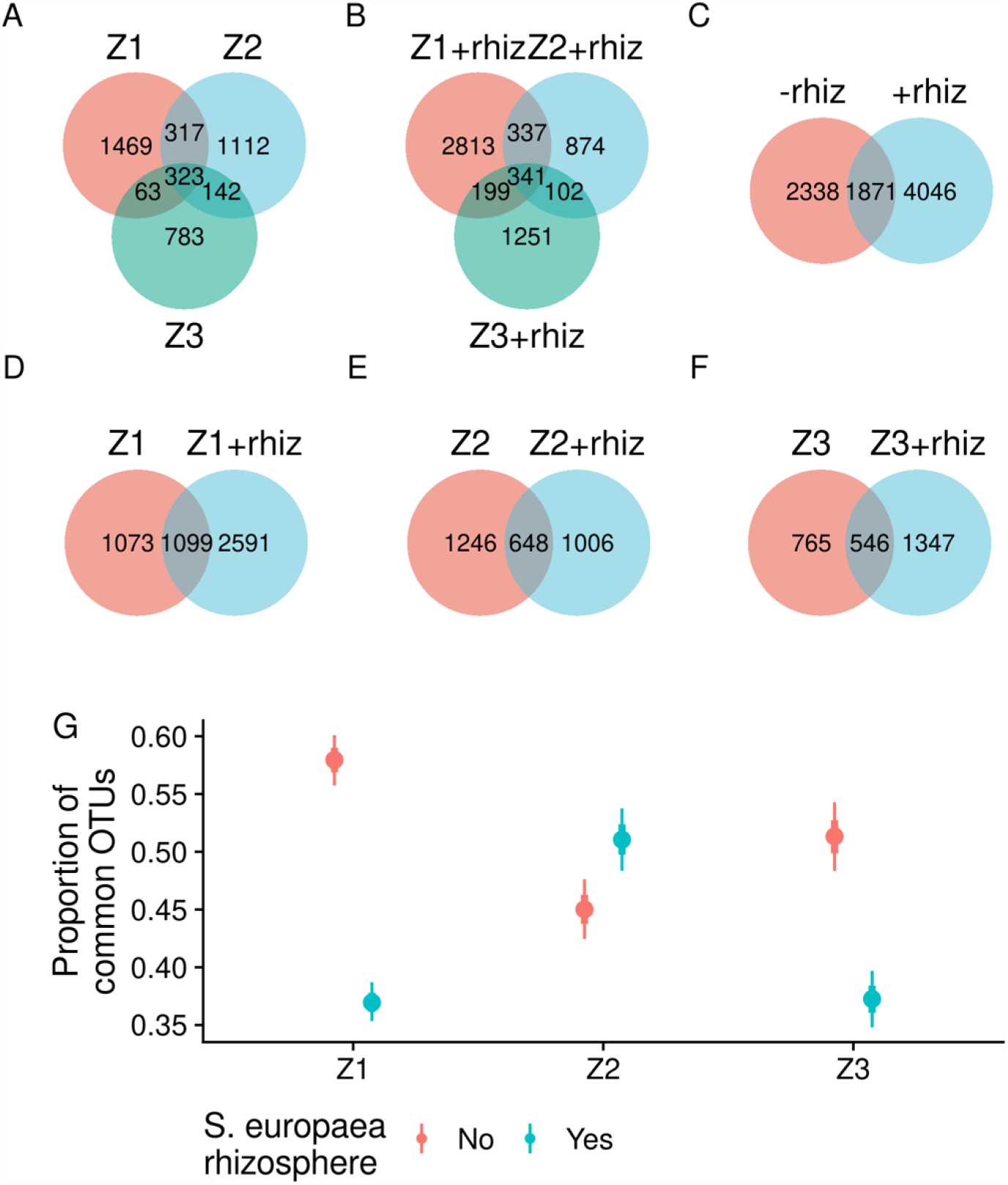
Relationships between OTU sets identified from *S. viminalis* roots grown in unsupplemented soils or in *S. europaea* rhizosphere-supplemented (rhiz) soils from three contamination zones. A. OTU sets from *S. viminalis* cuttings grown in unsupplemented soil samples from three zones. B. OTU sets from *S. viminalis* roots grown in *S. europaea* rhizosphere-supplemented soil from three zones. C. Relationship between OTU sets from *S. viminalis* cuttings grown in unsupplemented vs *S. europaea* rhizosphere -supplemented soils. D-F. Pairwise OTU sets of *S. viminalis* plants grown in unsupplemented or in *S. europaea* rhizosphere-supplemented soils from Z1 (D), Z2 (E), and Z3 (F). G. Proportion of common OTUs relative to total from *S. viminalis* cuttings grown in *S. europaea* rhizosphere-supplemented vs. unsupplemented soil from three zones. Points denote the best fit of the aggregated binomial model [common | trials(total) ∼ rhiz + zone + rhiz:zone]. Thick and thin lines denote 67% and 95% credible intervals, respectively. For OTU set comparisons, individual replicates were first rarefied and then merged by treatment groups.

The observed OTU richness and Shannon diversity index of bacterial communities in *S. viminalis* plants grown in unsupplemented soil were decreased in Z3 versus Z1 (effect sizes -413 OTUs, 95% CI, [33, -857], probability of effect size less than zero 0.975 and for Shannon index -1.1, 95% CI, [-0.46, -1.81], probability of effect size less than zero 0.999) and in Z3 versus Z2 (effect size -454 OTUs, 95% CI, [-18, -955], probability of effect size less than zero 0.977 and for Shannon index -1.0, 95% CI, [-0.32, -1.72], probability of effect size less than zero 0.996) (Fig 3 A, B). Whereas, there was no decrease in the inverse Simpson index across contamination zones in *S. viminalis* plants grown in unsupplemented soil (Fig 3 C). *S. europaea* rhizosphere treatments increased the observed OTU richness in Z1 and Z3, and the Shannon diversity index in Z3 (Fig 3 A, B).

**Fig 3.**
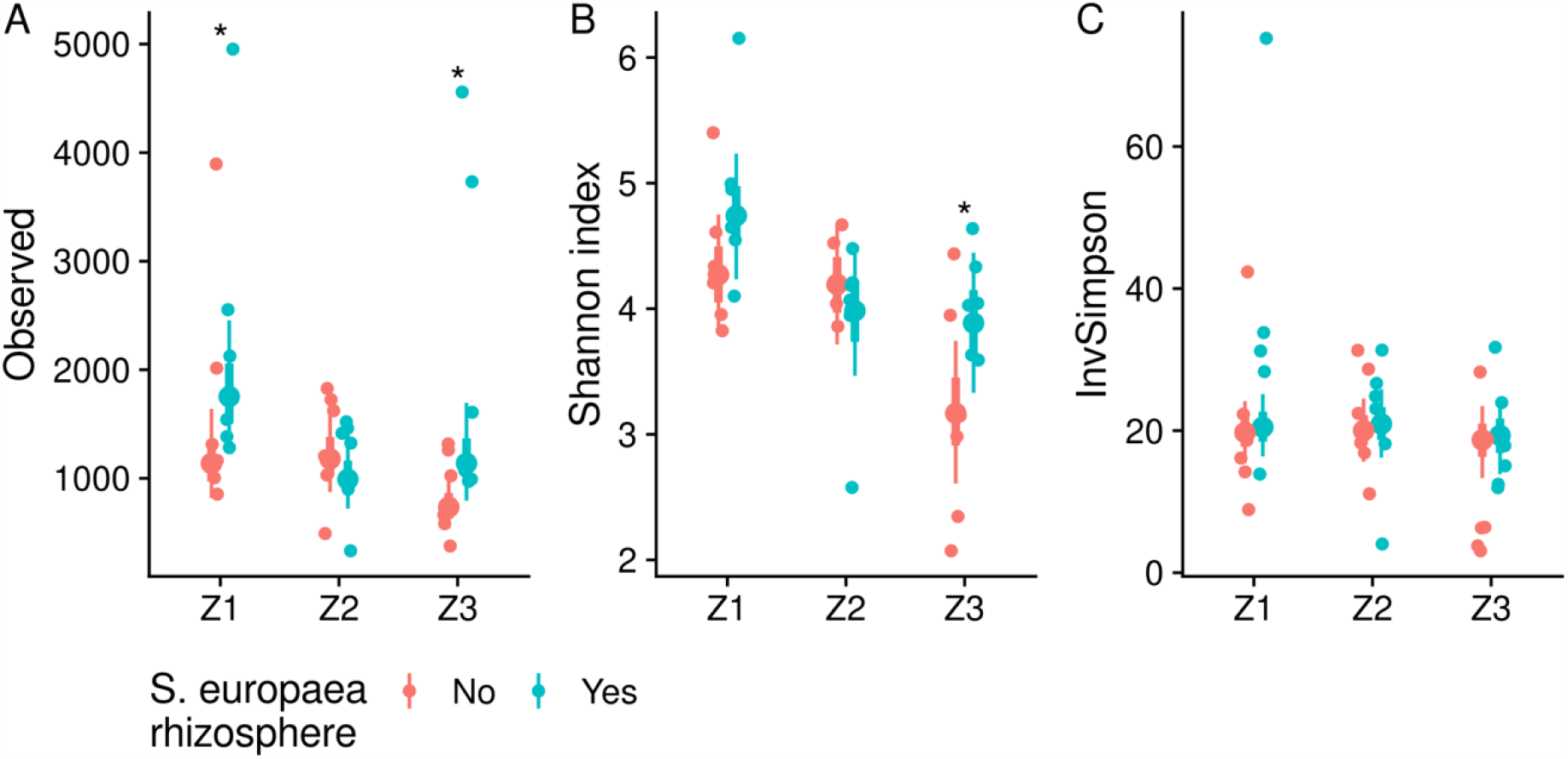
Alpha diversity measures of OTUs identified from *S. viminalis* roots grown in soils from three contamination zones with or without *S. europaea* rhizosphere supplementation. A. Conditional effects of the contamination zone and *S. europaea* rhizosphere supplementation to the observed OTU richness estimated from negative binomial model, adjusted for sequencing library size. B. Conditional effects of the contamination zone and *S. europaea* rhizosphere supplementation to the Shannon diversity index estimated from the robust linear model (Student’s t likelihood), adjusted for sequencing library size. G. Conditional effects of the contamination zone and *S. europaea* rhizosphere supplementation to the Inverse Simpson’s diversity index estimated from the robust linear model, controlled for sequencing library size. Non-rarefied OUT sets were used for diversity index calculations. Large points denote the model’s best fit. Thick- and thin lines denote 67% and 95% credible intervals, respectively. Small points denote individual observations, N = 6. Asterisks denote non-zero effect size in diversity measures (probability > 0.95) of *S. viminalis* cuttings grown in unsupplemented versus *S. europaea* rhizosphere-supplemented soils.

Beta diversity is illustrated in an NMDS plot based on Bray-Curtis distances of all samples (Fig 4 A). The samples clustered according to the pollution level along the NMDS1 axis with the more polluted clustering together in association positive NMDS scores. The NMDS plots also confirmed that communities from *Salicornia* rhizosphere-treated plants were clustering together with untreated communities from their zone of origin. There is some apparent overlap of some Z1 samples with Z2. The community dispersion was not significantly affected by the contamination zone or *Salicornia* rhizosphere treatment (Fig 4 B).

**Fig 4.**
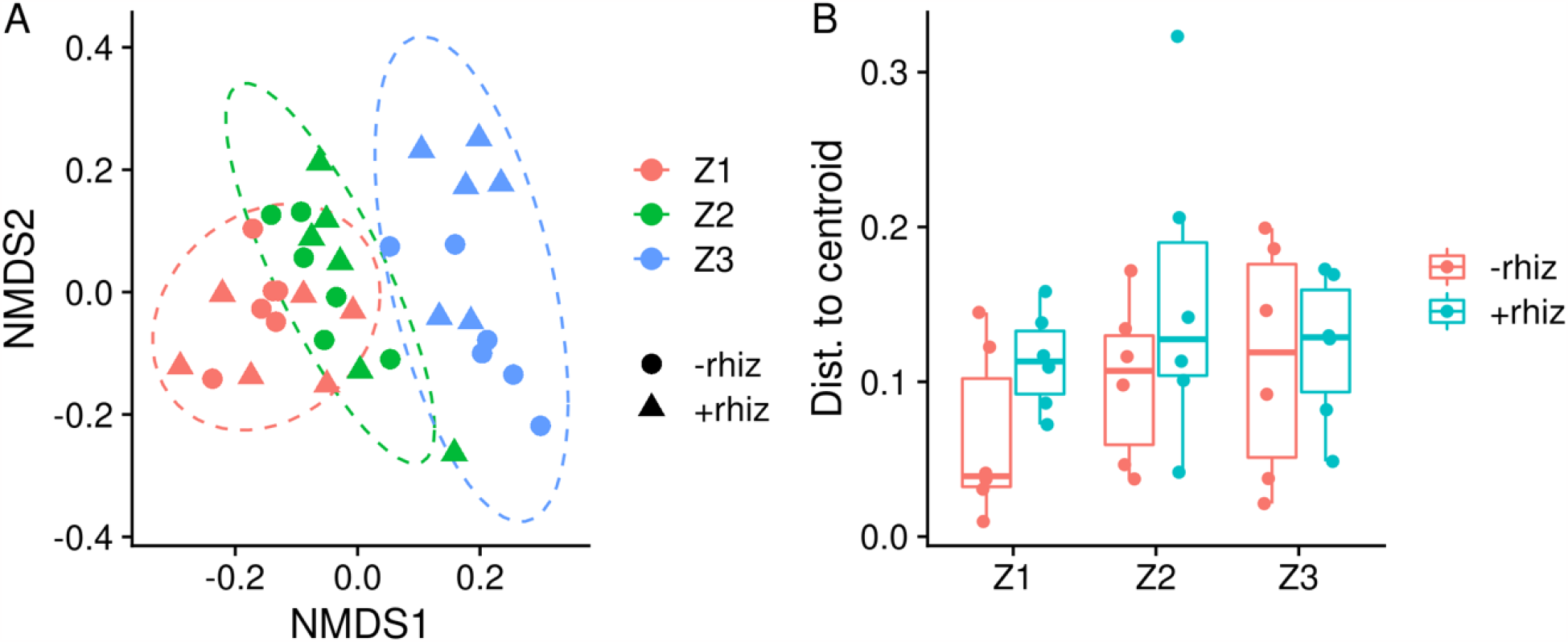
A. Non-metric multidimensional scaling (NMDS) using Bray-Curtis distances. Color denotes the contamination zone. The point shape denotes *S. europaea* rhizosphere supplementation. Ellipses are 95% confidence regions of t distributions. B. Distances to centroid calculated for NMDS Bray-Curtis distances.

Taxonomic analysis suggests a higher abundance of genera *Halomonas*, *Marinobacter*, *Idiomarina*, and *Marinimicrobium* in *S. viminalis* cuttings grown in Z3 soils (Fig 5).

**Fig 5.**
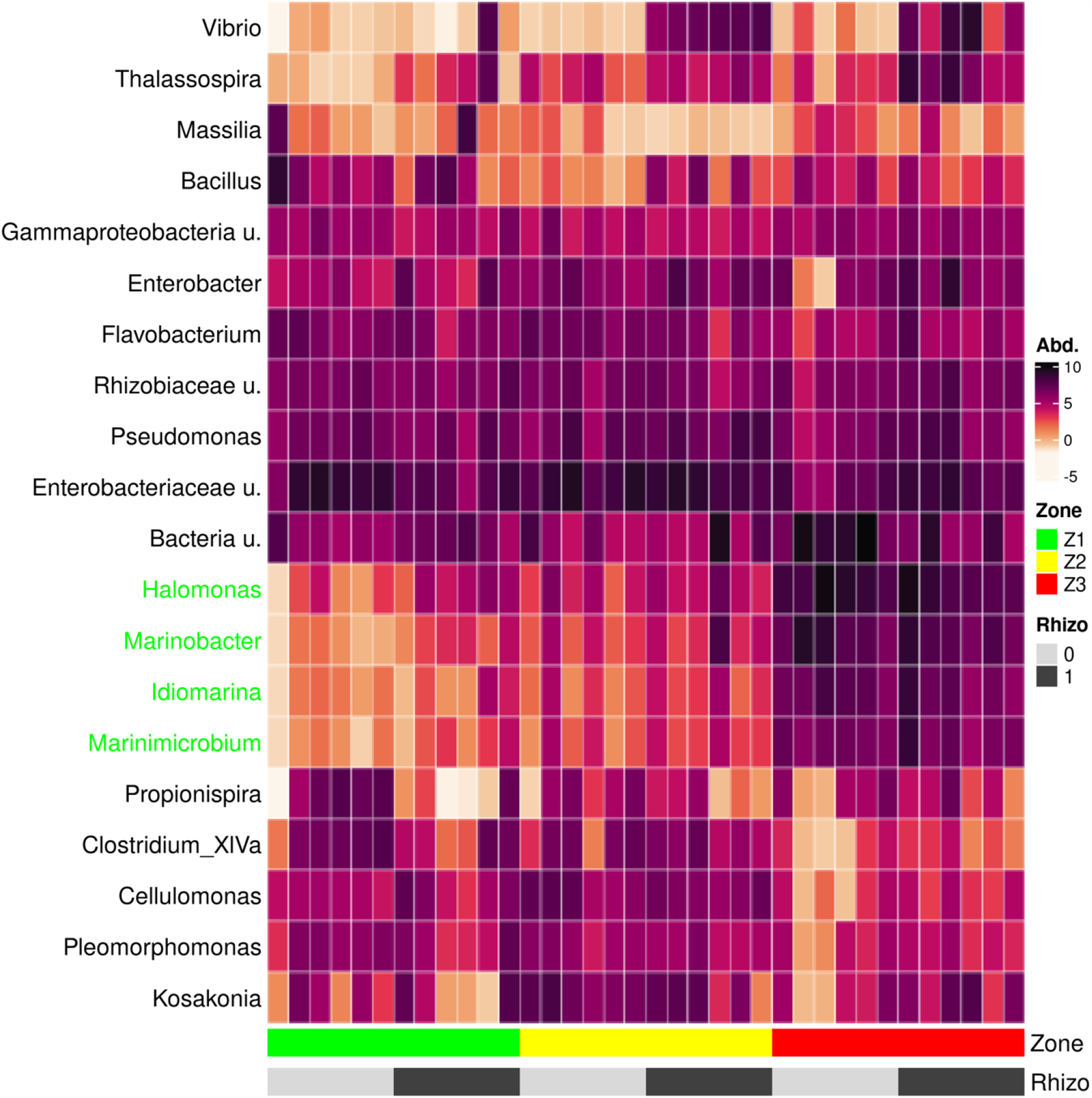
Relative abundance of the top twenty genera. Samples were ordered by the contamination zone and *S. europaea* rhizosphere supplementation.

## 4. Discussion

Some of the challenges that plants face in climate change are drought, flooding, salinity and heavy metal pollutants. Plant breeding has been generating the breeding lines capable of dealing with the stress situations. Considering, however, the rapid nature of environmental changes, it is unsure if the changes in the plant genomes are able to outcompete the challenges. Unlike the genes and regulatory regions of the genome, microbial composition can be rapidly modified by environmental cues, and may thus represent a mechanism for rapid acclimation and adaptations of individuals to a changing environment. Recently, we proposed the microbiota-mediated acclimatization concept, suggesting that changes in microbiota that originate from severe drought stress enable modern wheat cultivars to tolerate drought stress (Timmusk and Wagner 1999, Timmusk, Abd El-Daim et al. 2014, Timmusk and Zucca 2019, Timmusk, Pall et al. 2023). By the similar token here we hypothesize that the bacteria that have lived for long periods under HM and salt pollution stress have acquired the ability to tolerate the pollution; and are suitable for supporting bioremediation processes. The Stebnyk tailings storage long-term exposure to heavy metals and salts in the environment represents a threat to microbial populations, affecting communities and putting ecosystem integrity at risk (Fetsiukh, Bunio et al. 2022). The negative effects of salt pollution include changes in soil biogeochemistry and composition, and characteristics such as decreased aeration and decreased soil permeability with increased erosion. Our data representing the contamination gradient clearly showed that the alfa and beta diversity of the bacterial community decreased, and the community composition changed, along the pollution gradient, suggesting a homogenization of the communities as the pollution appears. The sequencing of the 16S rDNA library revealed the presence of a relatively rich endophytic community, indicated as the number of identified OTUs (Fig 1). The biodiversity and the richness of the bacterial endophytic community was assessed based on the observed number of OTUs, Shannon-, InvSimpson-, and evenness index (Fig 1 and 2). Modelling showed that the observed number of OTUs decreased approximately twice from zone 1 to zone 3, from 1408 (95%CI, 987-1999) to 678 (95%CI, 465-990), respectively (Fig 1). *S. europaea* supplementation significantly increased the number of OTUs in zones 2 and 3 (Fig 1). The biodiversity was significantly decreased across the contamination in un-supplemented plants with posterior probability >95% as indicated by the observed number of OTUs and Shannon indices (Fig 2). *S. europaea* supplementation lessened the decline based on observed OTUs, Shannon and evenness index (Fig 2). The inverse Simpson index was not significantly affected by the *S. europaea* treatments in any of the zones (Fig 2). This indicates that the number of dominating OTUs is not changing across zones in both *S. europaea*-treated, and untreated, bacterial populations (Fig 2), whereas decrease in richness is mainly caused by low-abundance OTUs.

While the pattern of Z1 and Z2 communities was relatively homogeneous, considerably larger changes in the community were observed in zone 3. *Marinobacterium*, *Idiomarina*, *Marinamicrobium*, and *Halomonas* were sparsely represented in communities Z1 and Z2 but were abundant in Z3. Many of the OTUs associated with the above-mentioned taxa have been frequently described as linked with *Marinobacterium*, *Idiomarina*, *Marinimicrobium*, and *Halomonas* representatives have earlier been chosen as inoculants to support growth under HM salt conditions (Desale, Patel et al. 2014, Mukherjee, Mitra et al. 2019). The rhizoremediation principle is that the plant roots colonize in the contaminated soil and associate with the subset of microorganisms present in the soil that help to tolerate the pollution stress. It may also involve metabolization of the pollutant. The endophytes are capable of degrading various organic pollutants and accelerating the extraction of toxic metals (Sonawane, Rai et al. 2022, Tufail, Iltaf et al. 2022). The bacterial metabolic pathways can synthesize natural chelators that improve the metallic availability for plant uptake and degradation (Saha, Tiwari et al. 2021, Haque, Srivastava et al. 2022, Raklami, Meddich et al. 2022, Sharma, Singh et al. 2022). It is important to understand that we can only speculate about the ecological role of the detected taxa in view of the fluid nature of bacterial taxa (Prosser, Bohannan et al. 2007, Timmusk, Pall et al. 2023) and even more so because the information on the taxa is based on what has previously been described in other systems. The basic biodiversity principle is that different organisms enhance the ecosystem functions under stress conditions. The observed increase in species richness can differentially influence the functioning of the ecosystem. If all species contribute approximately equally to the functioning of the ecosystem, the species effect may be additively decelerating if some of the species are to some extent functionally redundant. If the pool of species contains few species that can mediate handling pollutants efficiently, the effect will also be decelerating. Although several endophytes have shown bioremediation activities, microorganisms under field conditions face competition with a myriad of microbes naturally adapted to the environment. Hence, we identified natural endophytic strains of *S. viminalis* and *S. europaea* that would be capable of colonizing and efficiently supporting phyto-remediation based on the willows at the Stebnyk mine. How the microbial isolates behave under environmental conditions and what are the most influential consortia needs further study.

## In conclusion

Here we identified endophytic communities of the plant *Salix viminalis* that are affected by the pollution conditions. We show that *Salicornia europaea* supplements increase the abundance and species richness of the endophytic community. This, on the one hand, opens up new research avenues with implications for understanding complex interactions, as well as representing a promising source of novel microbial tools for the development of bioremediation biotechnology, the potential of which remains largely unexplored to date.

## Author contributions

ST conceived and designed the study; AF Stebnyk sampling, DNA extraction, PCR amplification, library preparation; TP processing the reads, taxonomic identification, statistical analysis; ST and TP wrote the manuscript; All authors contributed to the article and approved the submitted version.

## Acknowledgements

Dr. David Clapham is gratefully acknowledged for critically reading the manuscript. This study was supported by the Swedish Research Council VR 2017-05524 and VR2021-05471 to ST. Sequencing was performed by the SNP&SEQ Technology Platform in Uppsala. The facility is part of the National Genomics Infrastructure (NGI), Sweden, and the Science for Life Laboratory. TheSNP&SEQ Platform is supported by the Swedish Research Council and the Knut and Alice Wallenberg Foundation.

